# Status of Soil-transmitted helminthiasis among pregnant women attending antenatal clinic in Kilifi county hospital, Kenya

**DOI:** 10.1101/613570

**Authors:** Albert Njeru, Francis Mutuku, Simon Muriu

## Abstract

Soil-transmitted helminthes (STH) refers to parasites whose life cycle depends on a period of development outside the human host typically in a moist or warm soil. The most important geohelminths are; roundworms (*Ascaris lumbricoides*), whipworms (*Trichuris trichiura*) and hookworms (*Necator americanus* and *Ancylostoma duodenale*). The study aimed at investigating the current epidemiological status of soil-transmitted helminth infections among pregnant women in Kilifi County hospital (KCH) following the intervention of preventive chemotherapy.

Data collection and analysis involved evaluation of transmission trends of geohelminths from the previous surveys/ hospital records, use of direct fecal smears and Kato-Katz smears in estimation of prevalence and infection intensity and the use of structured questionnaires for the assessment of the predisposing factors associated with STH infections. Logistic regression analysis was employed to determine the risk factors associated with STH infections.

A total of 191 stool samples were collected and analyzed. Three species of soil-transmitted helminths (STHs) were identified with the overall prevalence of any STH infection being 16.75%. *Trichuris trichiura* was predominant (8.38 %), followed by *A. lumbricoides* (7.85%) then hookworms (6.29 %). About 10.99% of the participants had a single worm infection, 5.76 % had double co-infections and none of the participants had triple infection.

Preventive chemotherapy appears to give maximal returns in terms of reducing significantly the number of geohelminths reported since 2001. The study recommended the routine stool analysis in the antenatal profile, provision of alternative sources of iron to the pregnant women in order to reduce the tendency for soil consumption and the associated risk of STH infections, provision of safe water for domestic use, improved sanitation and proper personal hygiene (WASH).

## Introduction

Soil-transmitted helminth infections are among the Neglected Tropical Diseases (NTD) and remains a significant public health problem worldwide across the tropics and subtropics especially in many developing countries ^1^. Globally, more than 2 billion people or about 24% of the World’s population are infected with geo-helminths and over 300 million people suffer severe morbidity and even death ^2^. The main soil-transmitted helminths species affecting pregnant women are the roundworms (*Ascaris lumbricoides*),whipworms (*Trichuris trichiura*) and hookworms (*Ancylostoma duodenale* and *Necator americanus*).

Transmission of STH occurs when eggs passed in the faeces of infected people contaminate the soil in areas with inadequate sanitation ^3^. Mature helminths produce thousands of eggs in the intestines. Ingestion of eggs attached on raw vegetables (not washed or not peeled), contaminated water sources and eggs ingested by children who play in the contaminated soil eat without washing their hands are the factors that fuel the transmission of STH. Hookworm eggs hatch in the soil releasing larvae that mature and can penetrate the skin infecting people who walk barefoot on the contaminated soil. Direct person-to-person transmission or infection from fresh feces is not common because eggs passed in faeces need about 3 weeks to mature in the soil before being infective. Re-infection only occurs following contact with infective stages in the environment because soil-transmitted helminthes do not multiply within the human body ^3^.

The effects of STH infections during pregnancy include impairment of the nutritional status of the pregnant women through feeding on tissues and blood leading to iron and protein losses. Hookworms infections during pregnancy results to anaemia due to intestinal blood loss. STH infections also lead to increased nutrients malabsorption rates with roundworms competing for vitamin A in the intestines. Infected pregnant women usually have no appetite hence diminished nutritional intake and poor physical fitness. Trichiuriasis infections during pregnancy can cause diarrhoea and dysentery among pregnant women ^4^.

The burden associated with soil-transmitted helminth infections during pregnancy is related to the number of parasites harbored. Lightly infected pregnant women are usually asymptomatic while heavily infected pregnant women presents with wide range of symptoms including, abdominal pain, malnutrition, general malaise/ weakness, and impaired growth and physical development of the infant. High intensity STH infections leads to intestinal obstruction that requires surgical intervention ^5^.

In 2001,the World Health Organization endorsed a preventive chemotherapy intervention program aimed at reducing the morbidity associated with STH infections through periodic treatment of the vulnerable populations including pre-school children, school-age children and women of childbearing age (including pregnant women in the second and third trimesters and breastfeeding women) and adults in certain high-risk occupations ^6^. Adequate sanitation and health education reduces transmission and reinfection by encouraging healthy behaviors ^7^. Preventive chemotherapy during pregnancy is aimed at reducing the infection intensity and protecting pregnant women from the morbidity associated with STH infections ^6^.

The two drugs recommended for the treatment of STH infections in pregnancy are albendazole (400 mg) and mebendazole (500 mg) since they are effective, inexpensive(donated for free by the ministry of health), easy to administer and have minor side effects ^6^.

The aim of the current study was to investigate the status of soil-transmitted helminths infections among pregnant women seeking antenatal services at the Kilifi county hospital after fifteen years since the intervention of preventive chemotherapy targeting vulnerable populations. Undocumented reports indicate that since the year 2001, stool analysis for STH infections is not a routine in the antenatal profile among pregnant women attending antenatal clinics.

Pregnancy has been associated with increased prevalence of soil transmitted helminth infections compared to non-pregnant women ^8^. The epidemiology of Soil Transmitted Helminth infections, risk factors and the associated impacts during pregnancy is not clearly demonstrated although geohelminthiasis is common among many pregnant women residing in Kenyan rural areas ^9^. Stool analysis is not a routine in antenatal clinic profile since intervention of preventive chemotherapy in 2001 despite many pregnant women attending the antenatal clinic in Kilifi County Hospital not taking the albendazole and mebendazole prophylaxis yet 56 % practice geophagy according to previous study ^10^. The Ministry of Health through the Kilifi County government has made efforts to reduce the morbidity associated with STH infections among the vulnerable groups through the provision of preventive chemotherapy since the year 2001 according to the WHO guidelines but the impact is not documented.

Soil-transmitted helminth infections are among the most common infections worldwide and affect the poorest and most deprived communities with the greatest numbers reported in sub-Saharan Africa, America, China and East Asia ^11^. It is hoped that the results of this study will help consultants/researchers and policy operators in the health sector to understand the current epidemiological patterns of soil transmitted helminths and the preventive /control strategies employed together have on pregnant women and their fetus/newborns. Although the use of anti-helminthic prophylactic drugs alone has been shown to reduce morbidity and mortality due to soil transmitted helminths, it is not clear from the previous studies whether all pregnant women attending the antenatal clinics in Kenya adhere to taking the soil transmitted helminthic preventive chemotherapy and appliance of other control strategies like not eating soil as a form of iron supplement during pregnancy, good personal hygiene and adequate sanitation. It is also hoped that the results of this study will be adopted by the Ministry of Health (MoH), the county government of Kilifi and partners in formulating implementation strategies to curb soil transmitted helminths infections in the future in this region.

## Methodology

A cross-sectional hospital-based study was carried out among pregnant women who were attending the antenatal clinic at the Kilifi County hospital. Both primary and secondary data (from the previous hospital records/surveys and publications) were used. The population consisted of all the pregnant women who were visiting the Kilifi County hospital antenatal clinic. The population for recruitment in the study was all pregnant women who were seeking antenatal services at the Kilifi County Hospital. The subjects who admitted to have received anthelminthic drugs in 3 months prior to the study were excluded from the study.

### Data Collection

#### Evaluation of the transmission trends of soil-transmitted helminthes infections among pregnant women

The study reviewed the summaries of the monthly reported cases of each soil-transmitted helminth infection in the hospital records from the year 1996 (5 years’ pre-intervention of mass deworming which was started in 2001) and took a detailed look at studies that have been published in PubMed/google for the last fifteen (15) years relating to STH infections among pregnant women seeking antenatal services.

#### Estimating the prevalence of soil-transmitted helminthes infections among pregnant women

### Specimen Collection

A sample of fresh stool specimen was collected from all the participants. All the participants were provided with a labeled stool container, toilet paper, and applicator stick. Approximately 5 grams of stool specimens was collected into stool container using applicator sticks.

### Sample processing/Kato-Katz Technique

Each stool specimen was prepared using a sieve and put on a calibrated template for weighing in order to calculate eggs per gram of feces. The preparation on the glass slide was covered with a glycerin green impregnated cellophane paper. The preparation was turned upside down on a flat surface and pressed gently to spread the stool sample before examination. The slides were examined within one hour to avoid over clearing of hookworm eggs. All eggs in each preparation were counted to determine the number of eggs per gram of feces and all were classified as light infections^12^

### Assessment of the predisposing factors associated with soil transmitted helminthes infections

Data on socio-demographic factors, level of education, water access, sanitation, hygiene practices, geophagy, open defecation (presence of toilets/latrines), behavioral tendencies towards STH prevention/control source of soil and any history of deworming was collected from the participants using a pretested structured questionnaire during a face-to-face interview. A total of 191 questionnaires were administered to the 191 participants who provided the stool samples for STH infections analysis.

### Statistical analysis

All the data collected in the questionnaire and laboratory results was checked for completeness before entering it into Microsoft Office Excel spreadsheets. The data was analyzed using R-Statistical Package version 3.4.0. Logistic regression models were used to assess the association between independent variables considered and predisposing factors to STH infections. Soil transmitted helminthes infections were defined by the presence of any of the geohelminths species. The variables examined for their association with soil-transmitted helminthes infection (for both overall and by species as dependent variable) included; socio-economic status, level of education, source of water, household water treatment, marital status, age, gravidity, trimester of pregnancy, history of eating soil, source soil, information on open defecation and presence of pets in the house. Logistic regression was used to compute odds ratio and corresponding 95% confidence interval for those variables significantly associated with STHs prevalence. P-values less than 0.05 were considered to be statistically significant.

## Results

### Evaluation of transmission trends of soil-transmitted helminthes infections among pregnant women

A total of Five (5) yearly hospital records on STH infections among prenatal women were found for the period between 1996 and 2001 and no published journal was found for the five years prior to intervention of preventive chemotherapy. Between the year 2001 and 2016, a total 15 yearly hospital records and three (3) published studies relating to the 3 STH infections among pregnant women were reviewed while no study focused on either two or one of the STH species.

The figure below shows a summary of the transmission trends of the three STH infections among pregnant women seeking antenatal services in KCH since January 1996 to December 2016.A total of 907 cases of ascariasis infections were recorded and majority of the cases were reported in the year 1996 (148 cases) and the least were recorded in the year 2005 (4 cases). A total of 1090 cases were recorded for *T.trichiura* and majority of the cases were recorded in the year 1996 (237 cases) while the least (5 cases) occurred in the year 2004 as shown below. A total of 1338 cases of hookworm infections occurred and majority of the cases were recorded in the year 1996 (275 cases) and the least cases of hookworms infections were recorded in the year 2010 (7 cases) as shown below.

**Figure :**
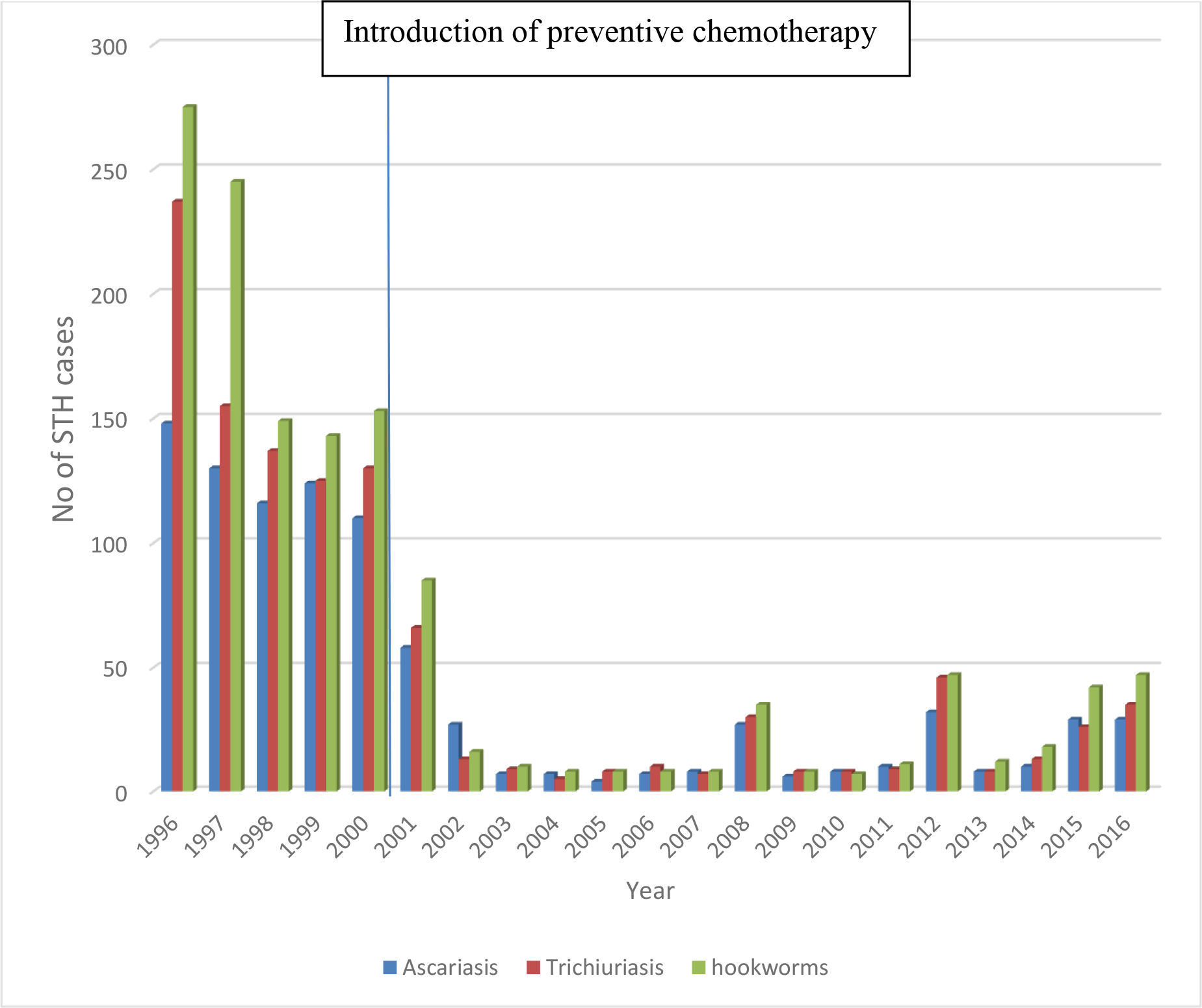
Trends of Soil-transmitted Helminthes infections among women attending the KCH antenatal clinic, 1996-2016.

### Estimating the prevalence of soil-transmitted helminthes infections among pregnant women

A total of 191 stool samples were collected and analyzed. Three species of soil-transmitted helminths (STHs) were identified in the stool samples with the overall prevalence of any STH infection being 16.75% (95% confidence interval (CI) 12.06-22.79%). *Trichuris trichiura* was the predominant intestinal helminth infection occurring in 8.38 % of pregnant women tested followed by *A. lumbricoides* (7.85%) and then hookworms (6.29 %). About 10.99% of the participants had a single worm infection, while 5.76 % had double co-infections and none of participant had all the three STH species considered as shown on table 1 below.

**Table 1:** Prevalence of soil-transmitted helminth infections among pregnant women attending antenatal clinic in Kilifi county hospital.

| **Soil-transmitted helminths** | **No. (%)** |
| --- | --- |
| *Ascaris lumbricoides* | 15 (7.85) |
| *Trichuris trichiura* | 16 (8.38) |
| *Necator americanus* | 10 (5.24) |
| *Ancylostoma duodenale* | 2 (1.05) |
| **Any soil-transmitted helminth infection** | **32 (16.75)** |
| Prevalence of single infection | 21 (10.99) |
| Prevalence of double infection | 11 (5.76) |

### Assessment of the predisposing factors associated with soil-transmitted helminthes infections

A total of 191 pregnant women attending antenatal clinic in Kilifi County Hospital were interviewed and provided stool samples for the analysis. Participants aged over 38 years were highly infected with soil transmitted helminthes with 10 out of the 13 assessed infected translating to a prevalence rate of 5.24% followed by the participants aged between 33-37 years 10 (5.24% infection rate) out of the 21 infected followed by participants aged 18-22 years who had a prevalence rate of 2.62% while persons aged between 23-27 years had the lowest infection rate of 1.05% (table 2).

**Table 2:**
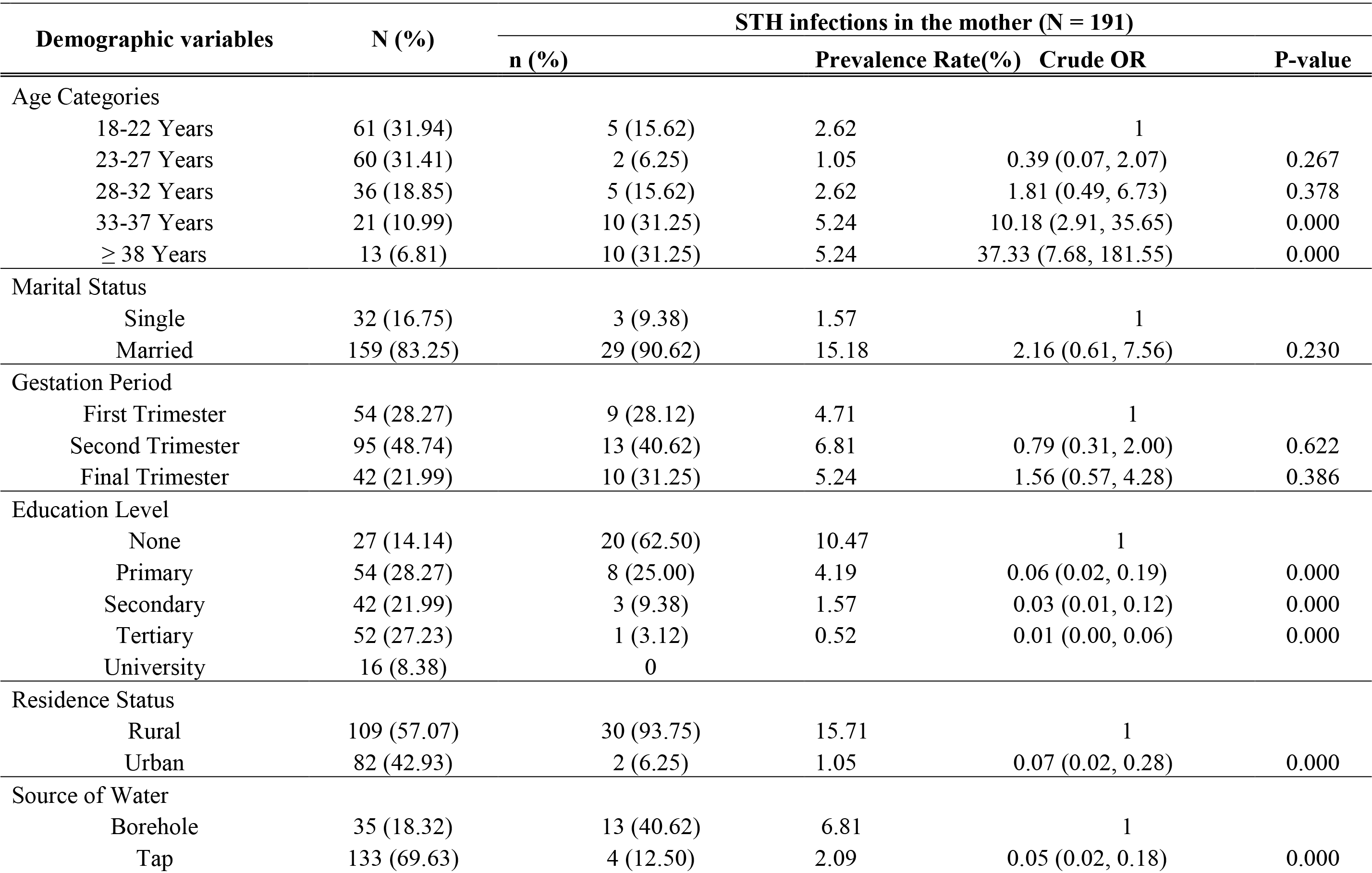
Univariate analysis of socio-demographic characteristics and their association with soil-transmitted helminths among women Univariate analysis of socio-demographic characteristics and their association with soil-transmitted helminths among women.

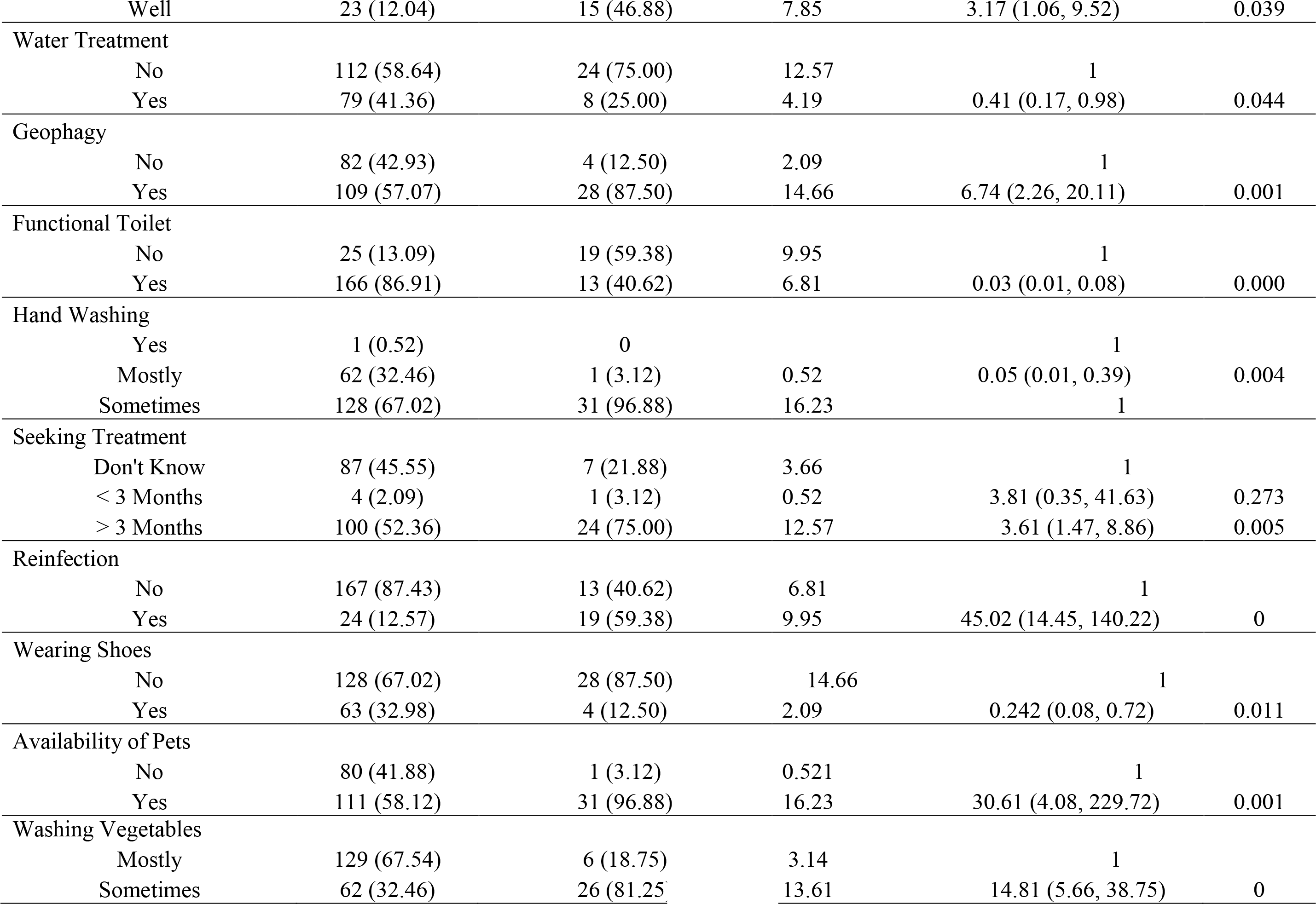

Majority of the respondents 83.29% (159) out of the 191 participants were married with 29 of them infected with STH which translated to an infection rate of 15.18% while 32 (16.75 %) of them were single and only three (3) of them were infected accounting for 1.57 % prevalence rate and none of the participants was either divorced or widowed or had any other marital status (table 2).

Majority of the respondents (57.07 %) lived in rural areas, out of whom 30 were infected with STH translating to an infection rate of 15.71% and 42.93 % (82) were from the urban areas of which two (2) of them were infected giving an infection (prevalence rate) of 1.05% in the urban area (table 2).

Respondents in the second trimester showed the highest infection rate of 6.81% followed by those in 3rd trimester who accounted for 5.24% infection rate and persons in the 1st trimester had the lowest prevalence rate of 4.71% (table 2).

The infections also varied according to educational background with those with no education reporting infection rate of 10.47%, primary 4.19%, secondary 1.57% and university/tertiary education which accounted for 0.52 % (table 2).

In terms of water sources for domestic purposes among the participants, majority of the infected respondents, 15(7.85%) used water from the wells followed by those who used borehole water, 13 (6.81%) and the least infected participants, 4(2.09%) used tap/piped water from Kilifi-Mariakani Water and Sewerage Company (KIMAWASCO) as shown on table 2. Out of the 191 participants interviewed, 8 out of the 79 who treated water were infected, hence infection rate among those who treated water before consumption was 4.19% while 24 out of the 112 participants who did not treat domestic water before consumption and the prevalence rate was 12.57% (table 2).

Geophagy was defined as the regular and deliberate consumption of non-food substances like soil. The study reported that 109 (57.07 %) study participants practiced geophagy while 82 (42.93 %) participants did not practice geophagy. 28 of the geophagic participants had STH infection rate of 14.66% compared to the 2.09 % (4) of the non-geophagic participants (table 2).

In relation to participants’ accessibility to functional toilets/pit latrines to determine probability of exposure to the geo-helminthes, 25 (13.09 %) had and no access to a functional toilet/pit latrine while 166(86.91%) of the participants had access to a functional toilet/pit latrine. Fewer of those with access 13(6.81%) had infections than those without access to toilet/latrines 19(9.95%) as shown on table 2.

Majority of the respondents 67.02% (128) washed their hands with soap/detergent sometimes before eating and after visiting the toilet and 31 of them were infected with STH infections which represented an infection rate of 16.23%. 62 (32.46%) of the respondents reported that they washed their hands with a soap/detergent mostly before eating and after visiting the toilet with only one (1) of them infected with STH which comprising 0.52% infection prevalence while one participant (0.52 %) reported to be washing her hands thoroughly with soap before eating and after visiting the toilet and she was not infected with any of the STH species (table 2).

129 (67.54%) participants mostly washed and cooked the vegetables properly before consumption and only six (6) of them were infected with STH infections which accounted for 3.14 % prevalence rate. However, the few women 62 (32.46 %) who washed and cooked their vegetables sometimes had a higher infection rate of 13.61 % (26 participants) as shown on table 2.

Histories of previous exposure/infection with STH infections of the respondents in order to establish the possible cases of re-infection were also reviewed. Majority of the respondents 167 (87.43 %) indicated that they had no previous history of infection with STH infections and only 13 of them were infected accounting for an infection rate of 6.81%. 24 (12.57%) of the participants reported to have had previous and frequent infections with STH with 19 of them infected representing a higher prevalence rate of 9.95% (table 2).

Majority of the respondents 52.36 % (100) had been de-wormed within a period of more than three (3) months prior to this study with 24 of them infected with STH which accounted for 12.57% infection rate while 45.55 % (87) didn’t know when they sought deworming and 7 of them were infected accounting for an infection rate of 3.66%.

However, 4 (2.09 %) of the respondents indicated that they had received the anti-helminth prophylaxis three (3) months prior to the study and only one respondent was infected with STH translating to 0.52 % prevalence rate (table 2).

Presence of pets in the homesteads and tendency to wear shoes was investigated in order to determine the chances of being exposed to geohelminths. Majority of the respondents 58.12% (111) had pets in their homesteads and 31 of them were infected with STH which accounted for 16.23 % while 80 (41.88 %) of the respondents did not have pets and only one subject was affected with STH which accounted for 0.52 % of the infections.

Majority of the respondents 128 (67.02 %) indicated that they did not wear shoes (walk barefooted/ wore slippers or open shoes) while at home with 28 of them infected who accounted for 14.66 % of the infections. 63 (32.98%) of the respondents reported that they never walk barefooted/wear slippers or open shoes while out of their houses and only four (4) of them were infected with STH infections accounting for 2.09 % infection rate (table 2).

There were significant associations between presence of any STH in the pregnant woman and geophagy (Odds Ratio (OR) 6.74; 95% CI 2.26-20.11., p = 0.001),taking water from the wells (OR 3.17, 95% CI 1.06-9.52, p = 0.039), history of re-infection (OR 45.02, 95% CI 14.45-140.22, p=0), taking anti-helmith prophylaxis after a period of more than three (3) months (OR 3.61; 95% CI 1.47-8.86, P=0.005), and availability of pets (OR 30.61; 95% CI 4.08-229.72) as shown on table 2.

There was a significant negative association found among those who treat water for domestic use (OR 0.41; 95% CI 0.17-0.98, p = 0.044) compared to those who don’t, those who wash hands mostly before eating and after visiting the toilet in relation to those who don’t, (OR 0.05; 95% CI 0.01-0.39, p=0.004), wearing shoes while at home in relation to those who don’t (OR 0.242; 95% CI 0.08-0.72, p=0.011),and those who use/have a functional toilet/pit latrine compared to those who don’t have/use (p for trend = 0.000 OR 0.03; 95% CI 0.01-0.08). In addition, a significant negative association was found in relation to urban living (OR 0.07; 95% CI 0.02-0.28, p = 0.000) as shown on table 2.

### Risk factors independently associated with STH infections among pregnant women attending antenatal clinic in KCH

Risk factors associated with STH infections included geophagy (adjusted OR = 14.640; 95% CI 2.922-73.362, p-value=0.001), vegetable washing (adjusted OR=4.964;95% CI 1.217-20.242, p-value=0.025),source of water (adjusted OR=5.837; 95% CI 0.036-1.124, p=0.036), presence of a functional toilet (adjusted OR = 14.640; 95% CI 0.046-0.853, p = 0.03),presence of pets and wearing shoes (adjusted OR=3.157;95% CI 0.325, 30.708, P=0.322) as shown on table 3 below.

**Table 3:** Multivariate analysis of independent risk factors for any soil-transmitted helminth infection in mothers attending ANC clinic at KCH.

| **Risk Factors** | **Adjusted Odds Ratio** | **95% CI** | **P Value** |
| --- | --- | --- | --- |
| Source of Water |  |  |  |
| Borehole | 1 |  |  |
| Tap | 0.290 | (0.121, 0.061) | 0.121 |
| Well | 5.837 | (0.036, 1.124) | 0.036 |
| Geophagy |  |  |  |
| No | 1.000 |  |  |
| Yes | 14.640 | (2.922, 73.362) | 0.001 |
| functional Toilet |  |  |  |
| No | 1.000 |  |  |
| Yes | 0.198 | (0.046, 0.853) | 0.03 |
| Pet in the Household |  |  |  |
| No | 1.000 |  |  |
| Yes | 3.157 | (0.325, 30.708) | 0.322 |
| Vegetable Washing |  |  |  |
| Mostly | 1 |  |  |
| Sometimes | 4.964 | (1.217, 20.242) | 0.025 |

## Discussion

This study demystifies the status of the three soil transmitted helminthe infections among pregnant women attending the antenatal clinic in Kilifi County Hospital since the intervention of the prophylactic antihelminth treatment in year 2001. The successful intervention of the anti-helminthic prophylaxis among pregnant women at the beginning of the year 2001 after a period of health education on prevention control measures led to significant reduction in the number of cases of STH infections reported compared to the period prior to the intervention of chemotherapy prophylaxis.

The current study revealed increased cases of the three STH infection species in 2008,2012,2015 and 2016 compared to other years which was associated with factors like negligence/poor/lack of proper health, stock outs, poor sanitation and poor hygiene. Mebendazole (500 milligrams per day for 3 days) or Albedazole (400 milligrams per day for 3 days) were the drugs of choice for the prophylaxis and management of STH cases which were mostly reported during the second and third trimesters of the pregnancy.

The respondents in the second trimester had the highest risk of the soil transmitted helminthe infections as compared to the other trimesters and this could be because women increasingly start practicing geophagy with increasing gestational age ^10^. The study revealed that women engage in geophagy during the first, second and third trimesters of pregnancy as found by other studies for instance the study by ^13^. In the present study, 57.07 % of the respondents practiced geophagy. The estimated prevalence of geophagy between and within countries is estimated between 10-75% ^14^.

The prevalence among pregnant women also ranged from 65% in Kenya, 46% in Ghana, 42% in Namibia to 28% in Tanzania. The results found on this study for the prevalence of geophagy were on the same limits as those found in coastal Kenya (56 %) but differed with those obtained in other studies done in the country, Central Kenya 26.1%, 45.7% in Bondo, Western Kenya and 74% in Nairobi, Kenya ^15^. The prevalence of STH infection observed in pregnant women was 16.75 %. The study showed that *Trichuris trichiura* had the highest prevalence of 8.8%, followed by *A. lumbricoides* with 7.85 % while hookworms had the lowest prevalence of 6.29 %. These study findings on the prevalence of STH infections were within the same range as obtained in another study carried out in Western Kenya by ^16^.

The study identified key predisposing factors associated with STH infection which are amenable for intervention like geophagy which accounted for 57.07% which concurred with the results of a study carried out in the same study region by ^10^, taking water from the wells, poor/lack of knowledge on water treatment, lack of hand wash before eating and after visiting the toilet, taking antihelmith prophylaxis after a period of more than three (3) months, and having pets and not wearing shoes while at home. The findings of this study provide some evidence for predictors to STH infections and public health prevention/control measures as reported by ^17^ who conducted a systematic review and meta-analysis of various studies published across the globe to examine the predisposing factors associated with the STH infections.

The findings of this study are unique in that they are from a cross-sectional hospital based study on pregnant women attending the antenatal clinic who are not routinely tested for STH infections upon enrolment but instead are given antihelminth treatment since the year 2001 as they are vulnerable to the infections. The study revealed the current epidemiological status and the public health burden of the soil transmitted helminthes among the pregnant women in Kilifi County along the Kenyan coast following the last fifteen years of selective deworming program.

### Limitations and challenges of the study

Stool analysis is not a routine in the antenatal profile therefore convincing mothers to bring stool was not easy therefore a single stool specimen was used to assess STH infection status which may underestimate the geo-helminth morbidity. Also the study did not compare the sensitivity of the Kato-katz technique with other diagnostic techniques like Polymerase Chain Reaction (PCR) due to feasibility issues.

### Conclusions and recommendations

STH infection is common among pregnant women attending antenatal clinic in Kilifi County Hospital. Risk factors associated with infection suggest that mass de-worming strategies must also address health education like discouraging pregnant women from geophagy, taking water from the wells, advocating on water treatment, hand washing before eating and after visiting the toilet, regular deworming after every three (3) months, and wearing shoes while at home.

The study recommended routine stool analysis for STH infections among the pregnant women attending the antenatal clinic, the provision of alternative sources of iron to the pregnant women in order to reduce the tendency for soil consumption and the associated risk of STH, supply of safe water for drinking and other domestic purposes, improved sanitation and proper personal hygiene (WASH).

### Ethical statement

Before commencing the study, the protocol was approved by the Pwani University Ethics Review Committee. Further approval was sought from the Kilifi County Hospital research committee. Written, signed or thumb-printed informed consents were also obtained from all participants. Soil-transmitted helminth cases were treated with Mebendazole.

## Acknowledgements

Alex Mutuku (KEMRI-Wellcome Trust) for his assistance and guidance during statistical analysis and all pregnant women who participated in the study.

